# Predator abundance attenuates positive density-dependent breeding spatial dispersion in a semi-colonial bird

**DOI:** 10.64898/2026.04.09.716047

**Authors:** Michał Wawrzynowicz, Lechosław Kuczyński

## Abstract

The spatial organisation of breeding populations can influence fitness and population dynamics, yet the spatial expression of density dependence may vary with ecological context. We investigated whether predator abundance alters this expression in the northern lapwing (*Vanellus vanellus*), a loosely social ground-nesting wader. Using long-term breeding-bird monitoring data from agricultural landscapes, we quantified spatial organisation using a Normalised Spatial Dispersion Index (NSDI) and modelled its relationships with lapwing density, hooded crow (*Corvus cornix*) abundance, habitat composition, and winter climate using generalised additive mixed models. Spatial organisation showed density dependence, but this relationship weakened progressively with increasing crow abundance. At low crow abundance, increasing lapwing density was associated with greater spatial dispersion, whereas the relationship approached zero at high crow abundance. Although it became slightly negative at the upper end of the crow-abundance gradient, uncertainty provided no clear evidence of a reversal to density-dependent aggregation. Broad-scale habitat composition and winter precipitation were not supported as predictors, whereas warmer winter temperatures were associated with greater aggregation. Our results show that predator context can attenuate the spatial expression of density dependence, suggesting that spatial organisation may be an overlooked component of variation in density-dependent population processes.

## Introduction

Density dependence is fundamental to population ecology because it describes how demographic and fitness-related processes change with population density [1,2]. Yet the strength and direction of these relationships need not be invariant across ecological contexts. Conspecifics can impose costs through competition while simultaneously providing benefits through cooperation, information transfer or reduced predation risk [3]. Ecological conditions that alter these costs and benefits may therefore modify not only the strength of the density dependence, but also its direction [4].

Spatial organisation is a particularly direct arena in which these opposing effects of density can be expressed [5]. Essentially, increasing conspecific density may intensify competition for breeding space and resources, favouring spatial dispersion [6]. Conversely, when individuals gain benefits from conspecifics’ spatial proximity, such as predator dilution, collective vigilance or communal defence, these costs can diminish and promote spatial aggregation [3]. Spatial organisation may therefore emerge from balance between density-dependent intraspecific interactions and external ecological factors, including predator presence and habitat quality [7].

Predation may be particularly important in shaping this balance. Experimental studies have shown that predation affects density-dependent prey dispersal [8,9], suggesting that the spatial consequences of increasing population density are contingent on predator context. Similarly, distinct predator communities have been shown to alter the strength and direction of density dependence in nest survival [4]. However, whether predation can similarly reverse the density dependence of breeding spatial organisation itself remains poorly understood.

We tested this possibility in the northern lapwing (*Vanellus vanellus*), a semi-colonial ground-nesting wader in which breeding arrangements range from solitary pairs to dense aggregations [10–12]. Lapwings maintain territories within aggregations [13] and engage in collective mobbing of nest predators [14,15], providing a system in which the costs and potential benefits of conspecific proximity can be highly context-dependent. Using long-term, large-scale breeding bird monitoring data, we quantified plot-level spatial organisation of breeding lapwings with a novel Normalised Spatial Dispersion Index (NSDI) and tested whether hooded crow (*Corvus cornix*) abundance modifies the relationship between local lapwing density and spatial dispersion.

We predicted that increasing hooded crow abundance, which acts as a proxy for predator pressure, will weaken the positive relationship between northern lapwing density and spatial dispersion. Specifically, under low hooded crow abundance, increasing northern lapwing density should increase spatial dispersion as competition for breeding space increases. Conversely, under high hooded crow abundance, the benefits of spatial proximity to conspecifics may offset competition costs, attenuating the density–dispersion relationship and potentially reversing its direction.

## Materials and methods

### a) Bird abundance data and spatial statistic

We obtained spatiotemporal abundance data for northern lapwing and hooded crow (2001-2024) from the Polish Common Breeding Bird Survey (CBBS), whose main methodological features follow those used in the British Breeding Bird Survey [16]. The scheme monitors fixed 1 km^2^ survey plots, each containing two parallel 1-km transects spaced 500 metres apart. Each transect is divided into five 200-m segments, and bird observation are recorded by segment and distance band from the transect line (“0–25 m”, “25–100 m”, “>100 m”, and “in flight”).

We treated ground observations recorded during the breeding season as indicators of breeding presence. Additionally, we only retained observations taken up to 100 metres from the transect line, and selected plots for the analysis, where at least three individuals were observed. Only observations covering the lapwing breeding season (from early April to mid-May) were retained. The final sample size was 606 data points form 238 survey plots (Figure 1a).

**Figure 1.**
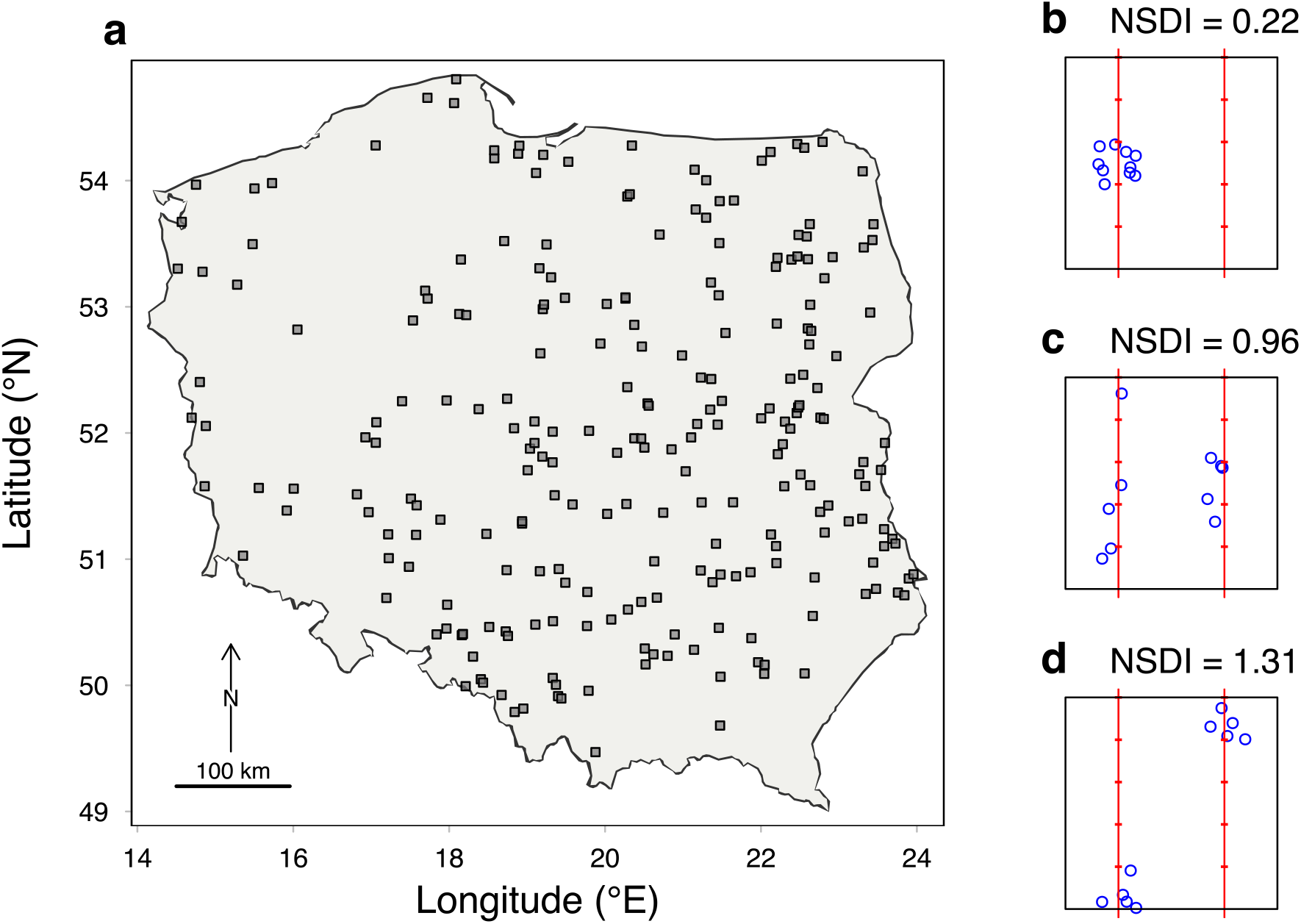
Quantifying breeding spatial organisation in northern lapwing. Panel **a** represents geographic distribution of the 606 breeding lapwing observations used in the analysis across 238 CBBS survey plots distributed across Poland. Panels **b–d** represent examples of reconstructed spatial configurations illustrating values of the Normalised Spatial Dispersion Index (NSDI): (**b**) 0.21, indicating spatial aggregation; (**c**) 0.96, indicating an approximately random configuration; and (**d**) 1.33, indicating spatial dispersion. Red lines indicate transects and the boundaries of the 200 × 200 m transect segments used for spatial reconstruction, whereas blue circles represent reconstructed spatial positions conditional on segment-level observations within the survey transect. Accordingly, NSDI, values <1 indicate greater aggregation than expected under complete spatial randomness (**b**), values ≈1 indicate random spatial organisation (**c**), and values >1 indicate greater dispersion (**d**).

To quantify plot-level spatial organisation of lapwing ground observations, we calculated Standard Distance Deviation from the survey plot geometric centroid. Because Standard Distance Deviation calculated on the raw counts is sensitive to abundance and survey plot geometry, it was normalised by the expected Standard Distance Deviation under a complete-spatial-randomness (CSR) null model (supplementary material, S2). Monte Carlo simulations indicated that Normalised Spatial Dispersion Index (NSDI) was not systematically related to abundance under the tested spatial configurations range (supplementary material, Figures S1, S2) and is interpreted as follows: values <1 represent spatial clustering, values ≈1 represent random spatial organisation, and >1 spatial dispersion (Figure 1b-c).

### b) Habitat and climatic data

To characterise temporal changes in habitat structure, we obtained annual land cover data from the Moderate Resolution Imaging Spectroradiometer (MODIS) Land Cover Type product (MCD12Q1 Version 6.1; Friedl & Sulla-Menashe, 2022). Bioclimatic variables were extracted from TerraClimate, which provides monthly global climate information with a spatial resolution of ~4 km [18]. Due to the high collinearity of monthly climate data, we aggregated them into 3-month seasonal variables (winter: December– February, by calculating their means (for temperatures) and logarithms of sums (for precipitation). Overall, we included four variables representing habitat composition and winter climatic conditions (Table S1) in our analysis.

### c)Statistical modelling

To analyse drivers of the plot-level spatial organisation of breeding lapwings, we used generalised additive mixed model (GAMM) [19] to allow for the flexible, non-linear modelling of the response curves. The Normalised Spatial Dispersion Index (NSDI) was modelled as the continuous response variable. Because the NSDI values are strictly positive and right-skewed (supplementary material, Figure S3), we assumed a Gamma distribution with a log-link function. To directly test our hypothesis that predation risk can attenuate the strength and reverse the sign of the density dependence of plot-level lapwing breeding spatial organisation, we incorporated a tensor product interaction (ti) between lapwing and crow abundances. Additionally, we included univariate smooths for conspecific density (log-transformed lapwing abundance) and predation risk (log-transformed crow abundance) to evaluate their independent effects, and landscape composition (the proportions of arable land and grasslands) and winter climatic conditions (minimum temperature and precipitation) to account for broad-scale environmental variation in spatial organisation. Finally, plot identity was included as a random intercept to account for repeated observations and persistent plot-level heterogeneity, while observer identity was included to account for observer-specific variation. Formally, the model was specified as follows:

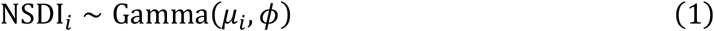

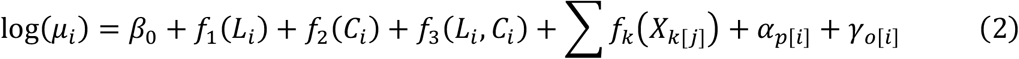

where NSDI_*i*_ is the calculated normalised dispersion index for survey *i*, with expected value *µ*_*i*_ and dispersion parameter *ϕ*. The term *β*_0_ represents the overall intercept. The non-linear smooth functions *f*_1_ and *f*_2_ denote the main effects of log-transformed lapwing (*L*_*i*_) and crow (*C*_*i*_) abundances, while *f*_3_ represents their tensor product interaction. The *f*_*k*_ functions represent smooth effects fitted to covariate vector *X*_*k*_ representing habitat and climatic covariates. Finally, *α*_*p*[*i*]_ and *γ*_*o* [*i*]_ are the independent, normally distributed random intercept effects for plot *p* and observer *o*, such that 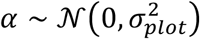 and 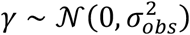.

All statistical analyses were performed in R version 4.5 [20].

## RESULTS

The full generalised additive mixed model (GAMM) explained 31.2% of the deviance in plot-level spatial organisation of breeding northern lapwings.

Lapwing density was positively associated with spatial dispersion (p < 0.0001), whereas the relationship with crow abundance was weaker but statistically supported (p = 0.0371). Importantly, the tensor product interaction between lapwing and hooded crow densities indicated that the density-dispersion relationship depends on crow abundance (p = 0.0442, F = 4.07). Because this interaction was close to the conventional significance threshold, we evaluated it using complementary sensitivity analyses. A leave-one-plot-out refitting analysis showed that the interaction was not dependent on individual survey plots (Figure S6), whereas a year-stratified permutation test yielded an empirical upper-tail probability p_perm_ = 0.0587 (95% Monte-Carlo confidence intervals: 0.0541, 0.0633; 9,999 permutations; Figure S7). Additionally, residual spatial autocorrelation was not detected across the observed spatial scales (Figure S5).

Habitat composition (arable land and grasslands) and winter precipitation were not supported as predictors of spatial dispersion, whereas minimum winter temperature showed a significant association (p = 0.0033; Table S2). The random-effects structure indicated significant among-plot heterogeneity (p = 0.0253), whereas observer identity explained little additional variation (p = 0.1948).

At zero crow abundance, increasing lapwing density was associated with greater spatial dispersion (Figure 2a). This positive density-dispersion relationship progressively weakened with increasing crow abundance (Figure 3). At the 95^th^ percentile of crow abundance, the estimated conditional effect of lapwing density was close to zero (*β* = −0.04, 95% pointwise CrI: −0.41 to 0.32), indicating substantial uncertainty around the sign of the relationship and providing no clear evidence for density-dependent aggregation.

**Figure 2.**
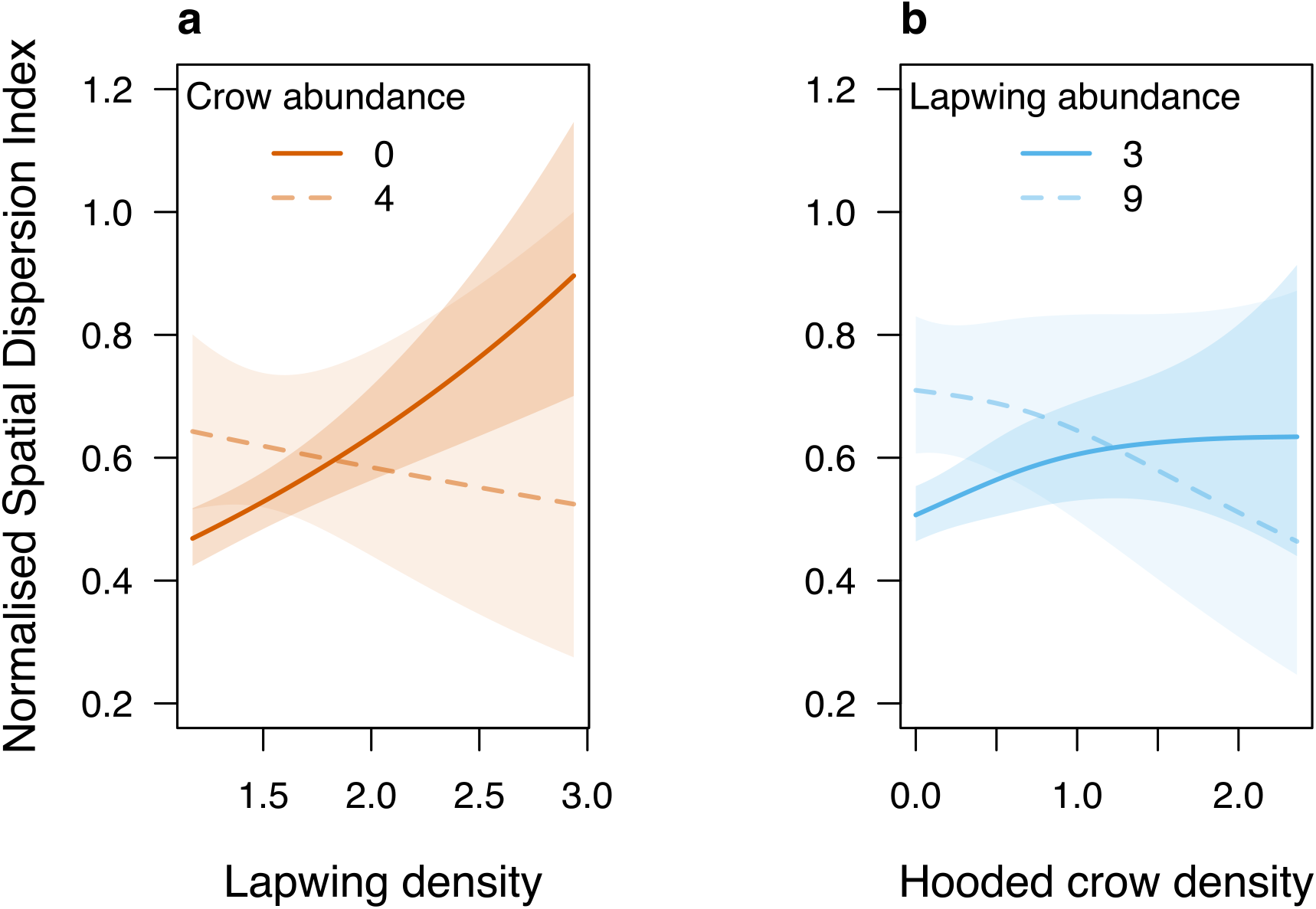
Predicted interaction effects between local conspecific density (log-transformed abundance) and local hooded crow density (log-transformed abundance) on the spatial dispersion (NSDI) of breeding lapwings. Estimates are derived from the tensor product interaction within the generalised additive mixed model (GAMM). Panels display the model-predicted NSDI (where <1 indicates spatial clustering, =1 indicates random distribution, and >1 indicates spatial dispersion) across the observed ranges of: (**a**) lapwing density [breeding pairs km^−2^] at two hooded crow abundance levels: 0 and 4 (≈95^th^ percentile); and (**b**) hooded crow density [breeding pairs km^−2^] at two observed lapwing abundances: typical (median) 3 and 9 (≈95^th^ percentile). Solid lines represent the fitted mean response each level, and shaded regions indicate 95% confidence intervals. Predictions isolate the interaction effect by holding all other continuous covariates at their median values and excluding random effects to show population-level trends.

**Figure 3.**
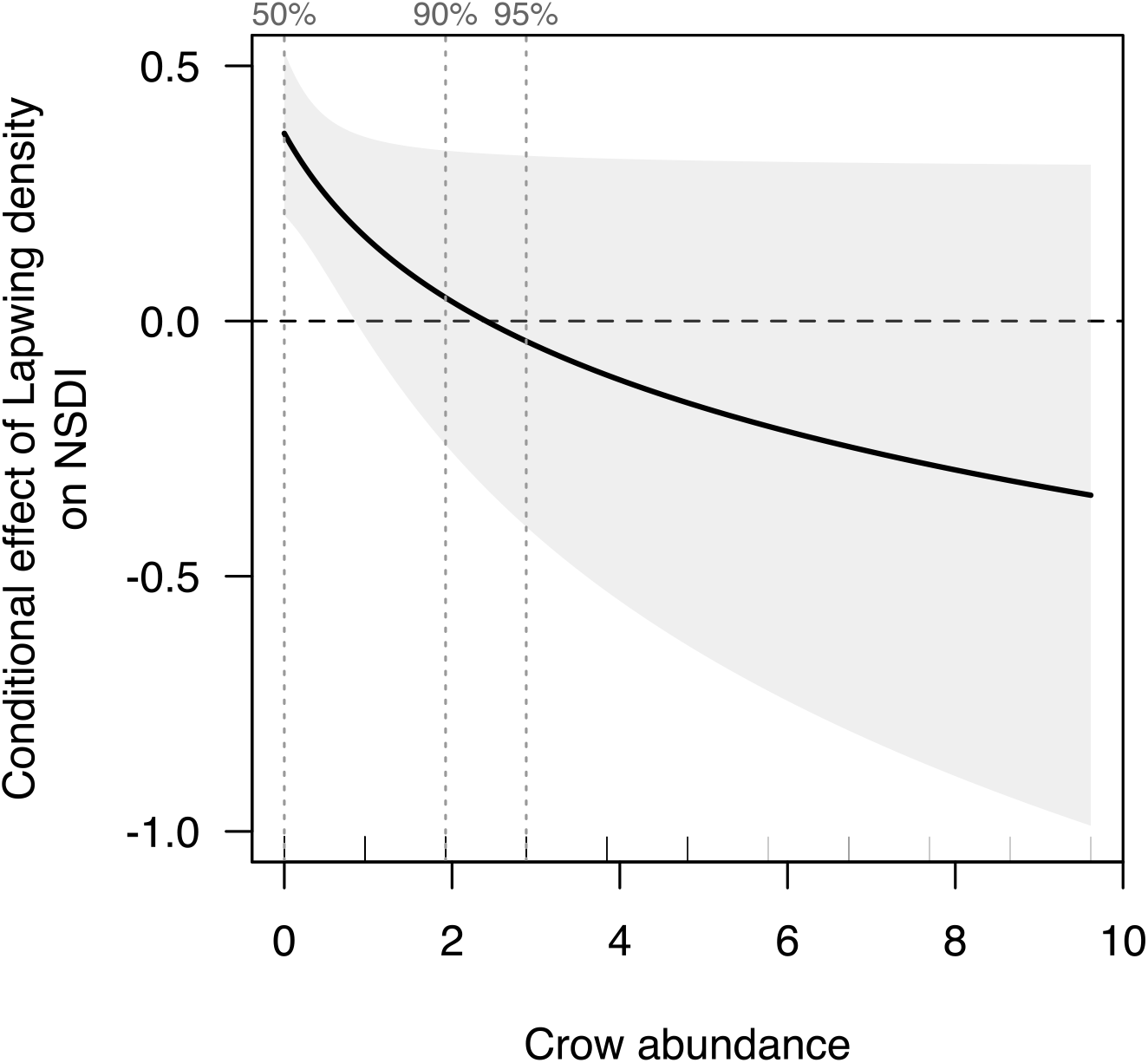
The conditional effect of lapwing density on spatial breeding dispersion (NSDI), estimated as the partial derivative of the log-transformed expected NSDI with respect to log-transformed lapwing abundance. The solid line represents the point estimate derived from the generalised additive mixed model (GAMM) tensor product interaction; the grey shaded polygon indicates the 95% pointwise credible interval. The horizontal dashed line denotes the zero-effect baseline (slope=0). Vertical dotted lines mark the 50^th^, 90^th^, and 95^th^ percentiles of observed crow abundance. Crow abundance is presented on the natural scale. Other continuous covariates are held constant at their sample medians, with random effects excluded from population-level inference.

## DISCUSSION

We found that the spatial expression of density dependence in breeding northern lapwings was contingent on hooded crow abundance. At low crow abundance, increasing lapwing density was associated with greater spatial dispersion, whereas this positive density–dispersion relationship progressively weakened with increasing crow abundance. Although the fitted relationship became slightly negative at the upper end of the crow-abundance gradient, uncertainty around the conditional effect remained substantial and overlapped with zero, providing no clear evidence for a reversal to density-dependent aggregation. Thus, local predator context attenuated the spatial expression of density dependence.

The positive density–dispersion relationship under low crow abundance is consistent with increasing competitive costs of conspecific proximity. Lapwings maintain breeding territories within aggregations and have a complex mating system [21–23], such that increasing local density can increase competition for breeding space, mates, and other resources, favouring greater spatial dispersion. At higher crow abundance, these costs may increasingly be offset by benefits of proximity to conspecifics, including collective vigilance, dilution and communal defence. However, our data do not distinguish among these mechanisms, nor do they demonstrate changes in individual spatial decisions; rather, they show that the spatial relationship between density and spatial dispersion changes systematically with predator abundance. This interpretation is consistent with experimental evidence that predator effects on spatial responses can depend on conspecific density [9], and extends this idea by showing that the density-dependent spatial organisation itself is context dependent.

We also found a significant association between warmer winter temperatures and greater spatial aggregation, whereas broad habitat composition and winter precipitation were not statistically supported. The temperature effect may reflect indirect changes in the availability or spatial distribution of suitable breeding habitat before breeding [24].

Nevertheless, this association was less central to our main interest and should therefore be interpreted as secondary. We hypothesise that under systematically warmer winter conditions, the availability of suitable, wet microhabitats required by lapwings for foraging becomes severely limited, effectively creating a landscape-scale bottleneck in breeding habitat availability. Under warmer conditions, contraction of suitable habitat could increase spatial aggregation by concentrating breeding birds into fewer suitable areas, consistent with evidence that lapwings can reduce territory size when suitable breeding resources are spatially concentrated [25,26]. Together, these associations suggest that spatial organisation can emerge from interacting biotic and abiotic constraints rather than from density dependence alone.

### a) Implications for density-dependent population processes

These findings may help to explain why density dependence can vary among populations and ecological contexts. Recently, Frauendorf et al. [4] showed that the strength and direction of density dependence in nest survival in semi-colonial oystercatcher (*Haematopus ostralegus*) varied among populations exposed to different predator communities. Whereas their analysis identified context dependence in the demographic consequences of density, ours identifies context dependence in the spatial organisation through which census density is expressed. Interestingly, these two processes need not be independent. If increasing density changes spatial organisation, and the direction or magnitude of this spatial response depends on predator abundance, then census density does not uniquely describe the social environment experienced by individuals.

This distinction is potentially important for understanding socially mediated Allee effects. Positive density dependence in fitness requires that increasing local density generates social benefits that outweigh density-dependent costs [27]. Yet the spatial realisation of density may itself be density dependent and predator-contingent, as indicated by our results. Consequently, the conditions under which social benefits can generate positive density dependence may vary across space and time rather than constitute a fixed property of a population. In this view, spatial organisation represents a potential intermediate process linking population density to fitness and, ultimately, to population dynamics.

This perspective also changes how density should be interpreted in spatially structured populations. For example, two sites can contain similar numbers of breeding individuals while presenting different local social environments because individuals use space differently. Conversely, density-dependent spatial reorganisation may alter the spatial scale over which social interactions operate. Thus, variation in density dependence may arise not only because the fitness consequences of density differ among environments, but also because density itself is expressed through different spatial configurations.

Our results also illustrate why the spatial scaling of social Allee effects may be more complex than population density alone suggests. Recent theory predicts that socially mediated demographic Allee effects depend on how individuals are connected across space and on whether local social conditions are maintained through the spatial distribution of individuals [28]. Our results suggest that this spatial realisation of density is itself context dependent, potentially altering the spatial scale over which social benefits accumulate. We therefore propose that variation in spatial organisation should be considered when evaluating how local component Allee effects may scale to population-level dynamics.

## 5 CONCLUSIONS

To conclude, we showed that the spatial expression of density dependence in breeding northern lapwings is contingent on predator context: increasing lapwing density was associated with greater spatial dispersion at low hooded crow abundance, but this association weakened as crow abundance increased. The result was robust to broad-scale spatial adjustment and exclusion of individual survey plots, while uncertainty at high crow abundance prevented us from resolving a reversal to density-dependent aggregation.

More broadly, our findings suggest that density should not always be treated as a sufficient description of the social environment within a population. The spatial configuration through which density is expressed may itself depend on the surrounding biotic and abiotic context, potentially contributing to heterogeneity in density-dependent demographic processes. Linking this spatial mechanism to fitness will require finer-resolution spatial data, direct measurements of predation rates, reproductive success, and behavioural experiments; however, our plot-level observational data cannot establish these mechanisms directly.

Spatial metrics such as our NSDI could nevertheless complement conventional population monitoring focused on abundance by quantifying how census density is spatially realised. Accounting for this spatial expression of density may improve interpretation of population responses to changing predator communities and environmental conditions.

## Acknowledgements

The authors would like to thank the MPPL volunteers who collected the data, the Chief Inspectorate for Environmental Protection (GIOŚ), and OTOP BirdLife Poland for providing the raw bird data. We would also like to thank Tomasz Chodkiewicz for his help with data curation and his valuable insights. Additionally, MW would like to thank Dominika Sikora for her insightful writing advice and overall support during the realisation of the study, and Adrian Wawrzynowicz for being silent co-founder of this study.

## Authors’ contributions

MW and LK conceived the ideas; LK and MW designed the methodology and prepared the data; MW and LK analysed the data. MW led the writing of the manuscript; LK acquired funding and provided overall supervision for the project. Both authors contributed critically to the drafts and gave final approval for publication.

## Data availability statement

This study uses novel code and third-party data. The code and processed data associated with the results are archived in a permanent Zenodo repository: https://doi.org/10.5281/zenodo.22280545. Raw bird population data used in the analysis are owned by OTOP BirdLife Poland (2000-2006) and the Chief Inspectorate for Environmental Protection (GIOSŚ, 2007-present). The authors do not have the rights to redistribute these raw data. OTOP data can be obtained upon request by contacting; GIOSx data are publicly available at: https://monitoringptakow.gios.gov.pl/PM-GIS/?lang=en.

