## Appendix S1 for "Predator abundance attenuates positive density-dependent breeding spatial dispersion in a semi-colonial bird"

### Electronic supplementary material S1: Land cover data

Table S1. Environmental and climatic variables selected for the analysis.

| Variable name | Description | Data source |
|---|---|---|
| Arable land | At least 60% of area is cultivated cropland [%] | MCD12Q1 Version 6.1 |
| Grassland | Dominated by herbaceous annuals (<2m) [%] | MCD12Q1 Version 6.1 |
| Tmin_winter | average of minimum temperature in December and from previous year and January and February from current year [°C] | TerraClimate |
| Precip_winter | logarithm of total precipitation in December from previous year and January and February from current year [mm] | TerraClimate |

To characterize temporal changes in habitat structure across Poland, we obtained annual land cover data from the Moderate Resolution Imaging Spectroradiometer (MODIS) Land Cover Type product (MCD12Q1 Version 6.1; (Friedl & Sulla-Menashe, 2022). This dataset provides global land cover classifications at 500-m spatial resolution derived from supervised classification of MODIS Terra and Aqua reflectance data. We selected the International Geosphere-Biosphere Programme (IGBP) classification scheme (Type 1), which categorizes land surface into 17 classes, including natural vegetation, agricultural systems, and urban areas (Sulla-Menashe et al., 2019).

Data granules covering the study area (tiles h18v03, h18v04, h19v03, h19v04) were acquired for the period 2001–2024 via the NASA AppEEARS platform. We processed the raw HDF files in R (R Core Team, 2025) using the terra package (Hijmans, 2026). For each year, granules were mosaicked using a modal rule to resolve overlapping pixel values and subsequently cropped to the extent of Poland.

To integrate categorical land cover information into our analytical grid, we calculated the fractional cover of each class within our sampling units. Binary masks were generated for each IGBP class and reprojected to the 1-km PUWG 1992 (EPSG 2180) reference grid using bilinear interpolation. This approach converted discrete pixel categories into continuous values ranging from 0 to 1, representing the proportional area coverage of each land cover type per grid cell. The resulting time series allowed us to quantify temporal variation in dominant habitat types, specifically focusing on grasslands and crops.

#### Electronic supplementary material S2: Reconstruction of spatial configurations, spatial statistic and its statistical properties

The MPPL survey protocol aggregates bird locations into 10 discrete spatial units per 1 km$^2$ plot. Each unit corresponds to a specific segment of the transect (200 m length) and includes all observations recorded within 100 m bands on either side of the transect line. Consequently, each bird count represents an individual located within a specific 200 × 200 m area (segment length × combined transect width).

To recover fine-scale spatial information from these aggregated counts, we applied an inverse mapping procedure. We treated the observation process as a discretization of continuous space; we reversed this by reassigning each observed individual to a coordinate drawn from a uniform distribution within the exact boundaries of the 200 × 200 m area where it was originally detected (segment centroid ± 100 m). This approach does not add arbitrary noise but rather restores the observations to their potential original locations, accounting for the spatial resolution lost during data collection.

We quantified the dispersion of these reconstructed point patterns using the Standard Distance Deviation ($SDD$), the spatial equivalent of standard deviation. $SDD$ measures the degree to which points are concentrated around their geometric centre and is calculated as:

$$\boldsymbol{SDD} = \sqrt{\frac{\sum_{i=1}^{n} \boldsymbol{d}_i^2}{\boldsymbol{n}}} \tag{1}$$

where $d_i$ is the Euclidean distance between the $i$-th individual and the mean center of the population, and $n$ is the total number of individuals in the plot.

Because the raw $SDD$ is sensitive to sample size ($n$) and the geometry of the plot, we normalized it against a null model of Complete Spatial Randomness (CSR). The $NSDI$ was defined as the ratio of the observed $SDD$ to the expected $SDD$ under random distribution:

$$\boldsymbol{NSDI} = \frac{\overline{\boldsymbol{SDD}}_{obs}}{\overline{\boldsymbol{SDD}}_{rand}} \tag{2}$$

To account for the uncertainty inherent in the spatial reconstruction and the stochastic nature of the null model, we used a Monte Carlo simulation approach ($R = 10{,}000$ iterations):

Observed ($\overline{SDD}_{obs}$): We generated $R$ spatial realizations of the observed count data using the reconstruction technique described above and calculated the mean $SDD$.

Random ($\overline{SDD}_{rand}$): We simulated $R$ random scenarios where the total number of observed individuals ($n$) was distributed across the 10 segments following a multinomial distribution with equal probabilities ($p = 0.1$). These random counts were then spatially reconstructed, and their mean $SDD$ calculated.

An $NSDI < 1$ indicates that the observed lapwings are more spatially aggregated than expected by chance (clustering), while an $NSDI > 1$ indicates spatial dispersion. An $NSDI \approx 1$ suggests a random spatial distribution. This normalization ensures that the index reflects active social aggregation or environmental constraints rather than artifacts of population density.

To ensure that our measure of spatial structure reflects spatial biological pattern rather than mathematical artifacts of population size, we tested the independence of the NSDI from local abundance. We performed Monte Carlo simulations across a range of biologically relevant abundances (n = 3 to 100 individuals per plot) under two boundary spatial configurations:

**Complete spatial randomness** (Figure S1): Individuals were distributed across segments according to a multinomial distribution with equal probabilities (p = 0.1).

**Complete clustering** (Figure S2): All n individuals were aggregated within a single, randomly selected segment (simulating extreme coloniality or habitat restriction).

For each abundance level (n), we calculated the NSDI using R = 10,000 iterations to stabilize the stochastic reconstruction component of the index. We assessed the relationship between abundance and NSDI values to verify the absence of systematic trend or bias.

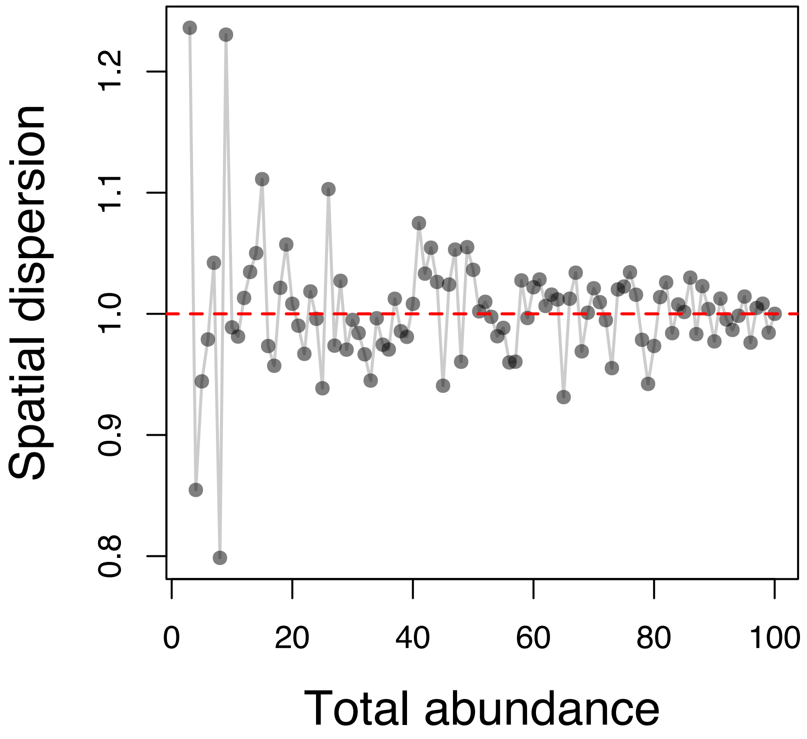


Figure S1. Assessment of the Normalised Spatial Dispersion Index (NSDI) under complete spatial randomness. The relationship between total plot abundance (n) and the calculated NSDI value when individuals are randomly distributed. The horizontal line indicates the expected value of 1.0. The results demonstrate that the index remains cantered on 1.0 across the range of abundances, confirming that the metric is unbiased with respect to local density. The variance decreases as n increases, consistent with the central limit theorem.

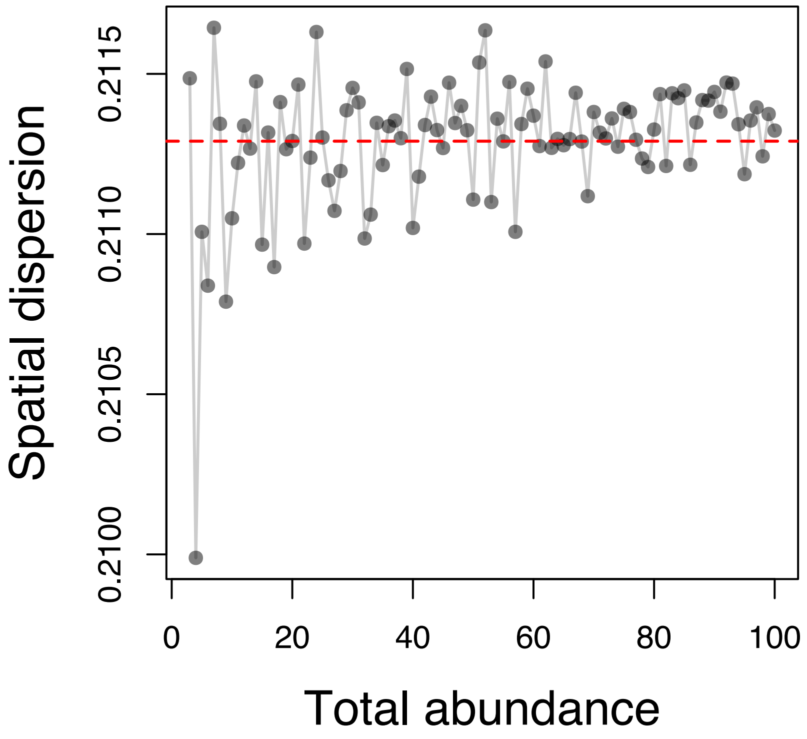


Figure S2. Assessment of the Normalised Spatial Dispersion Index (NSDI) under complete clustering. The relationship between total plot abundance (*n*) and the calculated NSDI value when all individuals are aggregated within a single spatial segment. The horizontal line represents the mean NSDI value across all simulations. The index remains constant on average regardless of the number of individuals in the cluster, indicating that the magnitude of the detected clustering is independent of the number of birds contributing to the cluster.

#### Electronic supplementary material S4: NSDI frequency distribution, modelling results and validation

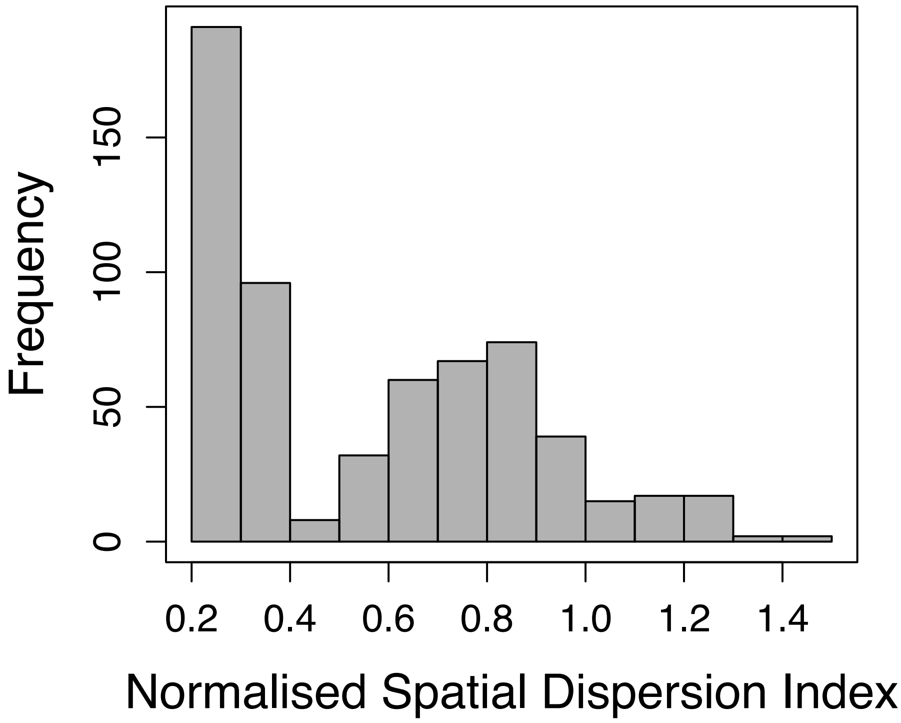


Figure S3. Frequency distribution of the Normalised Spatial Dispersion Index (NSDI) across all surveyed northern lapwing breeding plots. The histogram demonstrates the strictly positive, continuous, and strongly right-skewed nature of the spatial dispersion data. These inherent distributional properties justify modelling the response variable using a Gamma distribution with a log-link function in the generalised additive mixed models (GAMMs) used to evaluate the drivers of spatial organisation.

Table S2. Summary of the generalised additive mixed model (GAMM) evaluating the drivers of spatial breeding dispersion (Normalised Spatial Dispersion Index; NSDI) in the northern lapwing. The table presents the approximate significance of smooth terms, including main effects for conspecific density, predator abundance, habitat composition, and winter weather, as well as a tensor product interaction (`ti`) between conspecific and predator abundances. Random intercept effects were included for plot and observer identity. For each smooth term, the effective degrees of freedom (edf), reference degrees of freedom (Ref.df), F-statistic, and p-value are provided.

| *Parametric* | *Estimate* | *SE* | *t* | *p* |
|---|---|---|---|---|
| Intercept | -0.67 | 0.03 | -22.80 | **<.0001** |
| *Smooth terms* | *Edf* | *Ref. df* | *F* | *P* |
| Log lapwing density | 1.00 | 1.00 | 15.91 | **<.0001** |
| Log crow density | 1.60 | 1.93 | 3.00 | **0.0371** |
| Log lapwing density × Log crow density | 1.00 | 1.00 | 4.07 | **0.0442** |
| Arable land | 1.00 | 1.00 | 1.67 | 0.1969 |
| Grassland | 1.32 | 1.53 | 0.25 | 0.8196 |
| Winter precipitation | 1.00 | 1.00 | 2.91 | 0.0885 |
| Winter temperatures | 3.33 | 3.92 | 4.02 | **0.0033** |
| Observer ID (Random Effect) | 30.74 | 212.00 | 0.20 | 0.1948 |
| Plot ID (Random Effect) | 43.01 | 237.00 | 0.27 | **0.0253** |

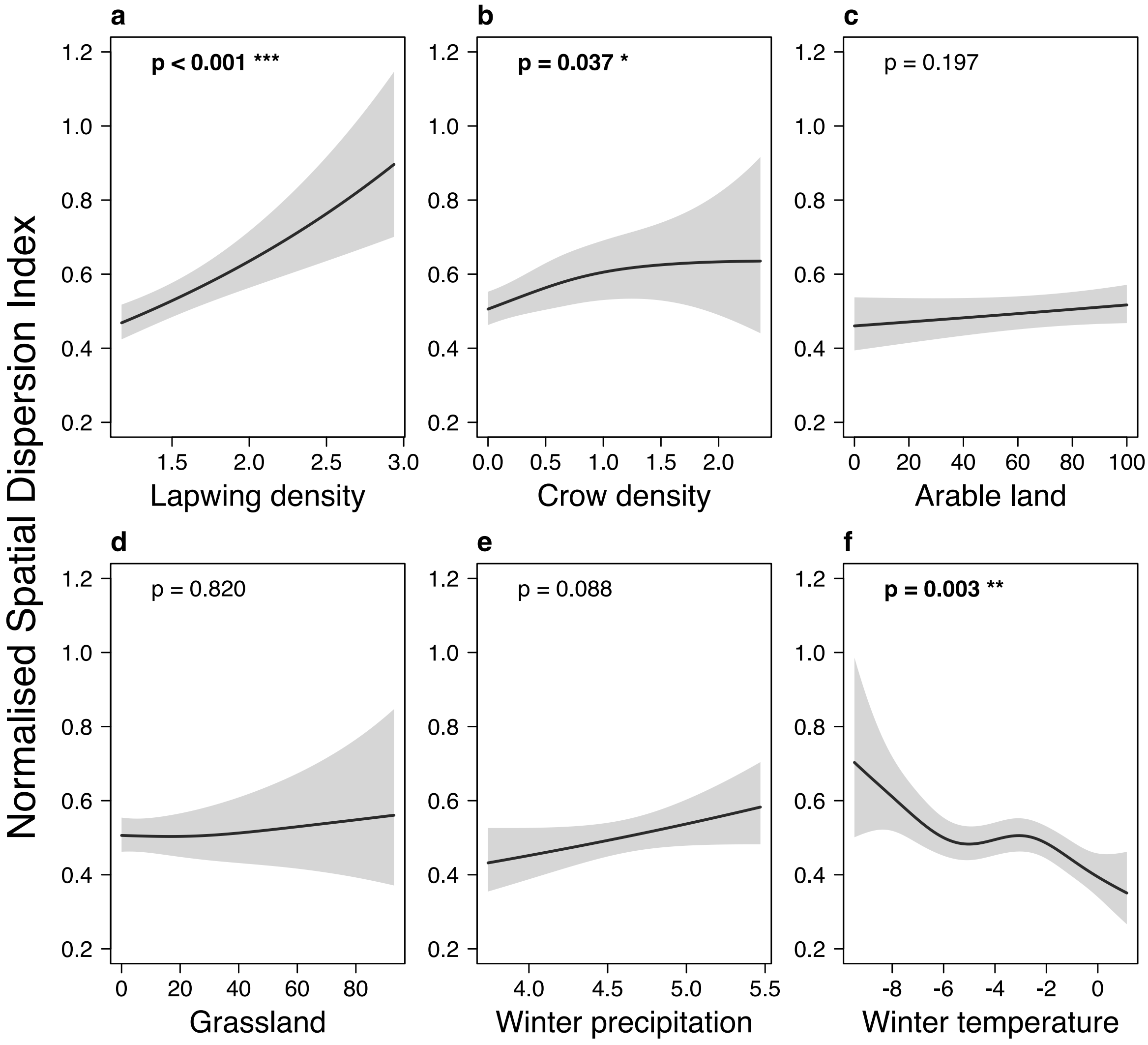


Figure S4. Marginal main effects of socio-ecological predictors on the spatial breeding dispersion of lapwings, quantified by the Normalised Spatial Dispersion Index (NSDI). Estimates are derived from the Generalized Additive Mixed Model (GAMM). The NSDI is interpreted as follows: values < 1 indicate spatial clustering, = 1 indicate random distribution, and > 1 indicate spatial dispersion. Panels illustrate the non-linear functional responses of spatial dispersion to: (a) local conspecific density [pairs $km^{-2}$]; (b) local predation risk (crow density [pairs $km^{-2}$]); (c, d) habitat composition (percentage cover of arable land and grasslands, respectively); and (e, f) pre-breeding winter weather conditions (log-transformed cumulative precipitation [mm] and average minimum temperature [°C] from December to February). Solid lines represent the fitted mean response, and shaded polygons represent 95% confidence intervals. To isolate main effects, non-focal continuous predictors were held constant at their median values.

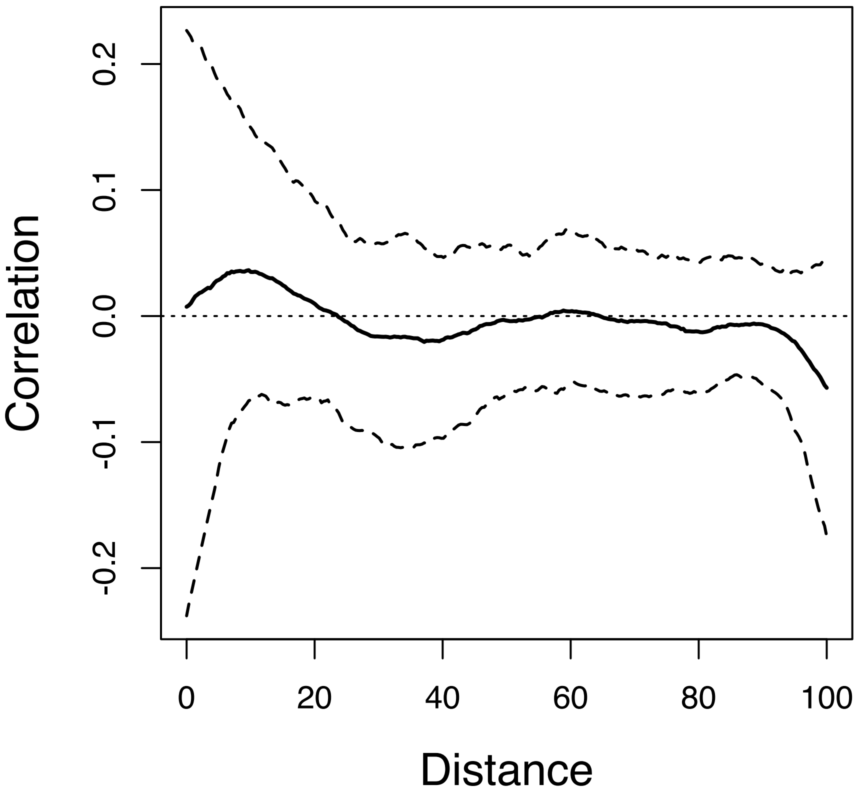


Figure S5. Spatial spline correlogram of the model residuals. The solid line represents the median spatial autocorrelation of the deviance residuals from the generalized additive mixed model (GAMM) across geographic distances. The dashed lines indicate the upper and lower 95% bootstrap confidence intervals, generated using 500 resampling iterations. The horizontal solid line at a correlation of zero indicates an expected lack of spatial autocorrelation. Because the 95% confidence intervals consistently encompass zero across the entire analysed distance spectrum, the correlogram showed no clear evidence of residual spatial autocorrelation across the analysed distance range.

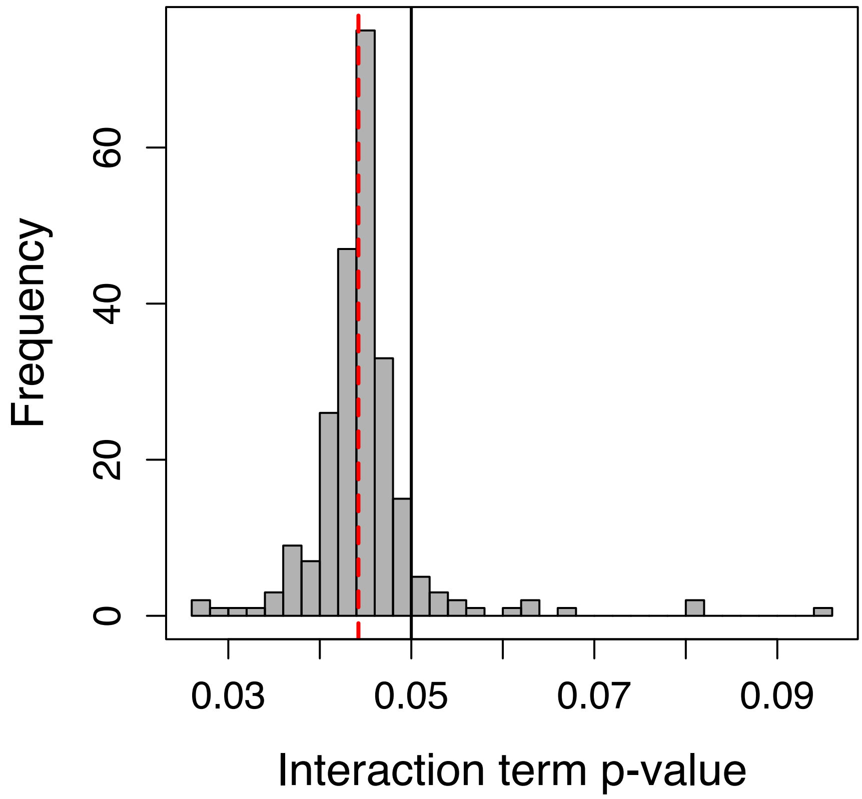


Figure S6. Frequency distribution of p-values for the interaction term derived from a leave-one-out sensitivity analysis. Each p-value represents the statistical significance of the tensor product interaction between local lapwing density and crow density when the generalized additive mixed model (GAMM) is refitted by iteratively excluding one study plot of 238 sampled plots at a time. The vertical solid line marks the conventional significance threshold of $p = 0.05$ and red dashed line represents p-value of the interaction term from the full model ($p = 0.0432$). The distribution demonstrates that the significance of the interaction term is robust to the exclusion of individual plots, as the leave-one-out p-values consistently cluster around the full-model estimate and largely (223 from 238) remain below the 0.05 threshold.

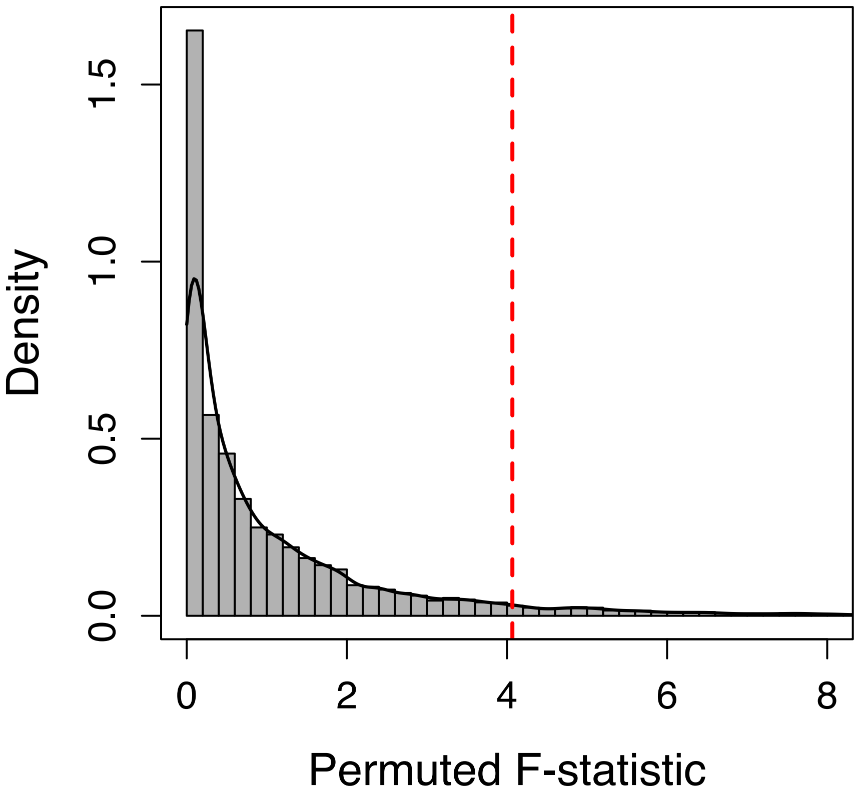


Figure S7. Empirical null distribution of the tensor product interaction term F-statistic obtained from 9999 year-stratified permutations, in which lapwing density was randomly reassigned among survey plot within years, and the GAMM was refitted. To make procedure computationally trackable, smoothing parameters where fixed at their estimates from the original model for each permutation; thus, the permutation test conditions on the smoothing parameters of the observed model. The vertical red dashed line represents the observed parametric F-statistic of the tensor product interaction term from the original GAMM (F = 4.07). The empirical upper-tail probability of obtaining a tensor product interaction F-statistic at least as large as observed was 0.0587 (Monte Carlo CI: 0.0541 and 0.0633).
